# p62/SQSTM1 selectively supports starvation-induced autophagy in N2a neuroblastoma cells

**DOI:** 10.64898/2026.01.03.697476

**Authors:** Arulventhan Marimuthu, Debasmita Saha, Varsha Vengataraghavan, Bangarupeta Murali Manpreet Jivin, Vasudharani Devanathan

**Affiliations:** Indian Institute of Science Education and Research, Tirupati, Andhra Pradesh, India

**Keywords:** Autophagy, p62 / SQSTM1, Autophagic flux, Starvation, Neuroblastoma cells, Bafilomycin A1, CRISPR–Cas9

## Abstract

Autophagy is a critical cellular process that maintains homeostasis and enables adaptation to metabolic stress. The selective autophagy receptor p62/SQSTM1 has been implicated in multiple aspects of autophagy regulation; however, its specific contribution to basal versus stress-induced autophagic flux remains incompletely defined, particularly in neuronal cells. In this study, we investigated the role of p62 in regulating basal and starvation-induced autophagy using a CRISPR–Cas9–generated p62 knockout (p62^−^/^−^) neuroblastoma (N2a) cell model. Autophagic flux was quantified by measuring LC3-II accumulation in the presence and absence of the lysosomal inhibitor bafilomycin A1. Under nutrient-replete conditions, basal autophagic flux was comparable between wild-type and p62-deficient cells, indicating that p62 is dispensable for constitutive autophagy. In contrast, starvation robustly increased autophagic flux in wild-type cells but failed to do so in p62^−^/^−^ cells, demonstrating a requirement for p62 in starvation-induced autophagy. Consistent with a stress-responsive role, p62 protein levels increased during starvation in wild-type cells under lysosomal inhibition, reflecting enhanced p62 expression rather than impaired degradation. Together, these findings reveal a context-dependent function for p62 in neuronal autophagy, whereby p62 is essential for mounting an effective autophagic response to nutrient deprivation but is not required for basal autophagic turnover.

## 1. Introduction

Autophagy is a conserved cellular process that maintains homeostasis by degrading damaged organelles, misfolded proteins, and toxic materials (Klionsky et al., 2016). It is especially vital in non-dividing cells like neurons, where impaired clearance can drive neurodegeneration (Nixon, 2013). The pathway involves forming double-membrane autophagosomes that capture cellular cargo and fuse with lysosomes for degradation, a process that can occur either broadly or with selective targeting of specific substrates (Mizushima et al., 2011).

Selective autophagy relies on adaptor proteins that link specific cargo to the autophagy machinery, ensuring the targeted removal of harmful aggregates and dysfunctional components. Among these adaptors, p62/SQSTM1 (Sequestosome1) is a key and evolutionarily conserved factor in metazoans (Komatsu, 2022). p62 is a 440–amino acid protein with multiple functional domains: an N-terminal Phox and Bem1 (PB1) domain, a ZZ-type zinc finger motif, and a C-terminal ubiquitin-associated (UBA) domain. These domains are bridged by an intrinsically disordered region (IDR) enriched with interaction motifs, including the LC3-interacting region (LIR), which connects p62 to autophagosomes, and the Keap1-interacting region (KIR), which integrates p62 into stress-response pathways (Komatsu, 2022). Functionally, p62 binds polyubiquitinated proteins, particularly aggregates of misfolded and ubiquitinated proteins, and is frequently observed in cytoplasmic inclusions across diverse protein aggregation disorders(Bjørkøy et al., 2005; Fan et al., 2010; Geisler et al., 2010; Ichimura et al., 2008; Komatsu et al., 2007; Pankiv et al., 2007). Beyond its role in cargo recognition, p62 also acts as a signaling hub, with its expression strongly upregulated by stress-responsive transcription factors such as NRF2, NF-κB, and MiT/TFE, underscoring its central role at the interface of autophagy and cellular stress adaptation(Katsuragi et al., 2015; Komatsu, 2022; Sánchez-Martin & Komatsu, 2018)

Recent findings reveal that p62 and related adaptors extend their role beyond cargo recognition to directly regulate autophagosome biogenesis. In selective autophagy, receptors such as p62, TAX1BP1, and OPTINEURIN interact with FIP200 of the ULK1 complex and Atg9 to initiate isolation membrane formation around targeted (Kageyama et al., 2021; Nakamura et al., 2020; Ravenhill et al., 2019; Turco et al., 2019; Vargas et al., 2019). Subsequent binding to Atg8–PE, together with wetting effects, drives elongation of the isolation membrane along the cargo surface (Agudo-Canalejo et al., 2021; Kageyama et al., 2021). Thus, adaptor proteins such as p62 act not only as cargo selectors but also as active contributors to autophagosome formation.

In the present study, we examined the turnover of p62 under basal conditions in N2a cells, a widely used neuronal precursor–like model, and found that p62 turnover was largely preserved under nutrient-replete conditions. This observation suggests that p62 is not essential for maintaining basal autophagic flux in this neuronal cell model. To further define the role of p62 in autophagy regulation under metabolic stress, we generated a p62 knockout (p62^-/-^) N2a cell line using CRISPR–Cas9–mediated gene editing. This model enabled systematic investigation of the contribution of p62 to basal and starvation-induced autophagy.

## 2. Materials and Methods

### 2.1 Cell Culture

Mouse neuroblastoma N2a cells (ATCC) were cultured in Dulbecco’s Modified Eagle Medium (DMEM, cat. no. 11995065, Gibco, USA) supplemented with 10% fetal bovine serum (FBS, cat. no. 16000044, Gibco, USA) and 1% penicillin-streptomycin (cat. no. 15140122, Gibco, USA). Cells were maintained at 37°C in a humidified atmosphere with 5% CO2. Cells with passage 5 were used for all CRISPR/Cas9 experiments.

### 2.2. sgRNA Design and Vector Construction

The genomic sequence of the mouse SQSTM1 gene was retrieved from the Ensembl genome database. Single guide RNAs (sgRNAs) targeting exon 1 of SQSTM1 were designed using multiple computational tools including E-CRISP, CHOPCHOP, and CRISPOR to ensure optimal targeting efficiency and minimize off-target effects. Two sgRNAs were selected based on their predicted cutting efficiency and specificity scores.

The sgRNAs were cloned into the pU6-(BbsI)_CBh-Cas9-T2A-mCherry vector (Addgene plasmid #64324) for simultaneous expression of Cas9 nuclease and mCherry fluorescent protein.

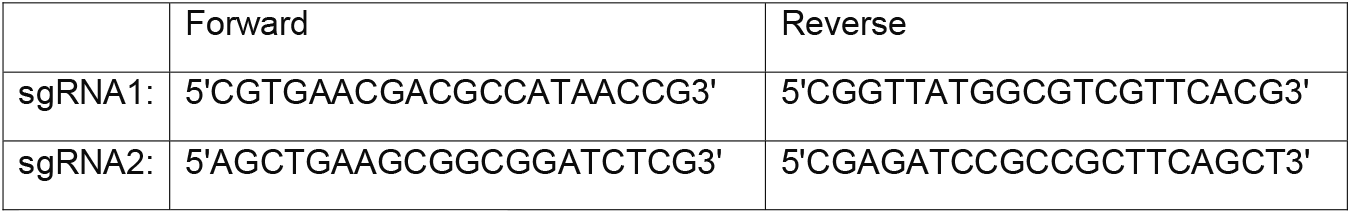

### 2.3 CRISPR/Cas9 Transfection

N2a cells were seeded in 6-well plates and grown to approximately 70-80% confluence. Transfection was performed using Lipofectamine 3000 reagent (cat. no. L3000001, Thermo Fisher Scientific) according to the manufacturer’s recommended protocol. Briefly, cells were transfected with the sgRNA-expressing plasmids in regular culture medium. After 24 hours, the transfection medium was replaced with fresh complete medium, and mCherry expression was analyzed 48 hours post-transfection using fluorescence microscopy.

### 2.4 Single Cell Cloning and Selection

Transfected N2a cells were harvested by trypsinization and diluted to achieve single-cell seeding density in 96-well plates using limiting dilution cloning. After 24 hours of plating, individual wells were examined for mCherry fluorescence to identify successfully transfected clones. mCherry-positive single cell clones were sequentially expanded through 24-well, 12-well, and 6-well plates following confluence at each stage. Upon reaching confluence in 6-well plates, each clone was split into two 6-well plates: one for continued propagation and maintenance, and the other for genomic DNA extraction and protein expression analysis.

### 2.5 Knockout Validation

Genomic DNA was extracted from selected clones and subjected to whole genome sequencing analysis performed by Eurofins Genomics to confirm successful knockout of the SQSTM1 gene. Additionally, protein expression analysis was conducted to verify the absence of p62/SQSTM1 protein in knockout clones compared to wild-type controls.

### 2.6 Optimization of drug Bafilomycin A1 Treatment Conditions

N2a cells were seeded in complete DMEM (4.5 g/L glucose, 10% FBS, 1% penicillin-streptomycin) and incubated for 24 h before drug treatment. Cells were then exposed to Bafilomycin A1 at 5, 10, 20, 40, or 80 µM for both 2 hours and 4 hours. Also, DMSO was used as the vehicle control for all treatment conditions. Following treatment, cells were washed with cold 1× PBS and processed for lysate preparation. LC3-II levels were quantified by western blot to identify the minimal effective Bafilomycin A1 dose and treatment time required to inhibit lysosomal degradation. The optimized dose was subsequently applied in all downstream autophagy assays.

### 2.7 Protein Lysate preparation

Following the indicated treatments, culture medium was aspirated and cells were washed once with ice-cold 1× PBS. Cells were lysed in 1× RIPA buffer (Sigma, 20-188) supplemented with 0.1% SDS, a protease inhibitor cocktail (1×; Roche Life Science, Germany; 11873580001), and a phosphatase inhibitor cocktail (1×; Roche Life Science, Germany; 04906837001). Lysates were collected into microcentrifuge tubes and incubated on ice for 30 min with intermittent mixing. Insoluble material was removed by centrifugation at 16,000 × g for 30 min at 4 °C. The clarified supernatants were collected and used for protein quantification and subsequent Western blot analysis.

### 2.8 Assessment of Autophagy Under Basal, Starved, and Pertussis Toxin–Treated Conditions

Neuro2a wild-type (WT) and Sqstm1/p62 knockout (KO) cells were seeded and maintained in DMEM media for 48 h prior to experimental manipulation unless otherwise indicated. To assess basal autophagy, the cells were treated with Bafilomycin A1 (10 nM) for 2 h. Next, to induce starvation-mediated autophagy, culture medium was replaced with Earle’s Balanced Salt Solution (EBSS) 4 hours before protein lysates. Cells were treated with Bafilomycin A1 (10 nM) for 2 hours before harvesting. DMSO was used as the vehicle control. Following treatment, cells were immediately processed for protein extraction.

### 2.6 Western blotting

Equal amounts of protein (20 µg per lane) were mixed with 1× Laemmli sample buffer containing 5% β-mercaptoethanol and boiled at 95 °C for 5 min. Proteins were resolved by SDS–PAGE on 14% polyacrylamide gels using a running voltage of 80 V for the stacking gel and 120 V for the resolving gel for approximately 2 h. Proteins were transferred onto 0.2 µm PVDF membranes by wet transfer at 100 V for 90 min. PVDF membranes were activated in methanol prior to transfer. Membranes were blocked in 5% BSA prepared in TBS-T (20 mM Tris–HCl, 150 mM NaCl, 0.1% Tween-20) for 1 h at room temperature and incubated overnight at 4 °C with the following primary antibodies: rabbit anti-LC3B (Cell Signaling Technology, #3868; 1:2000), mouse anti-p62/SQSTM1 (Abcam, ab56416; 1:2000), and mouse anti-β-actin (Abcam, mAbcam 8226; 1:5000). After washing, membranes were incubated with HRP-conjugated anti-rabbit IgG (Cell Signaling Technology, #7074; 1:10,000) or anti-mouse IgG (Cell Signaling Technology, #7076; 1:10,000) secondary antibodies for 1 h at room temperature. Immunoreactive bands were detected using Clarity Western ECL substrate (Bio-Rad, 1705060). Membranes were incubated with substrate for 2–3 min prior to imaging and developed at multiple exposure times to ensure non-saturating signal acquisition. Densitometric analysis was performed using Fiji (ImageJ). LC3-II band intensities were quantified and normalized to β-actin. Data represent three independent biological replicates.

### 2.6 Statistical analysis

Statistical analyses were performed using GraphPad Prism (version 10.6.1). Comparisons between two groups were conducted using unpaired two-tailed Student’s *t*-tests with Welch’s correction, as data were derived from independent samples and equal variance was not assumed. Autophagic flux was calculated as the difference between LC3-II levels measured in the presence and absence of bafilomycin A1. Individual data points represent biological replicates. A *p* value < 0.05 was considered statistically significant.

## 3. Results

### 3.1 Generation of CRISPR–Cas9–mediated p62/SQSTM1 knockout cell line

To investigate the role of p62/SQSTM1 in neuronal autophagy, we generated a p62-deficient N2a cell line using CRISPR–Cas9–mediated genome editing. N2a cells were transfected with a Cas9–sgRNA expression vector containing an mCherry reporter to facilitate identification of transfected cells. Robust mCherry fluorescence was detected 48 h post-transfection, indicating efficient delivery of the CRISPR construct (Fig. 1A). Bright-field imaging showed no overt changes in cell morphology following transfection. Single-cell–derived clones were isolated by limiting dilution and expanded sequentially. Multiple clones (designated B, C, F, and G) were successfully propagated and maintained stable growth characteristics during expansion (Fig. 1B). Based on growth behavior and transfection marker expression, these clones were selected for molecular validation.

**Figure 1.**
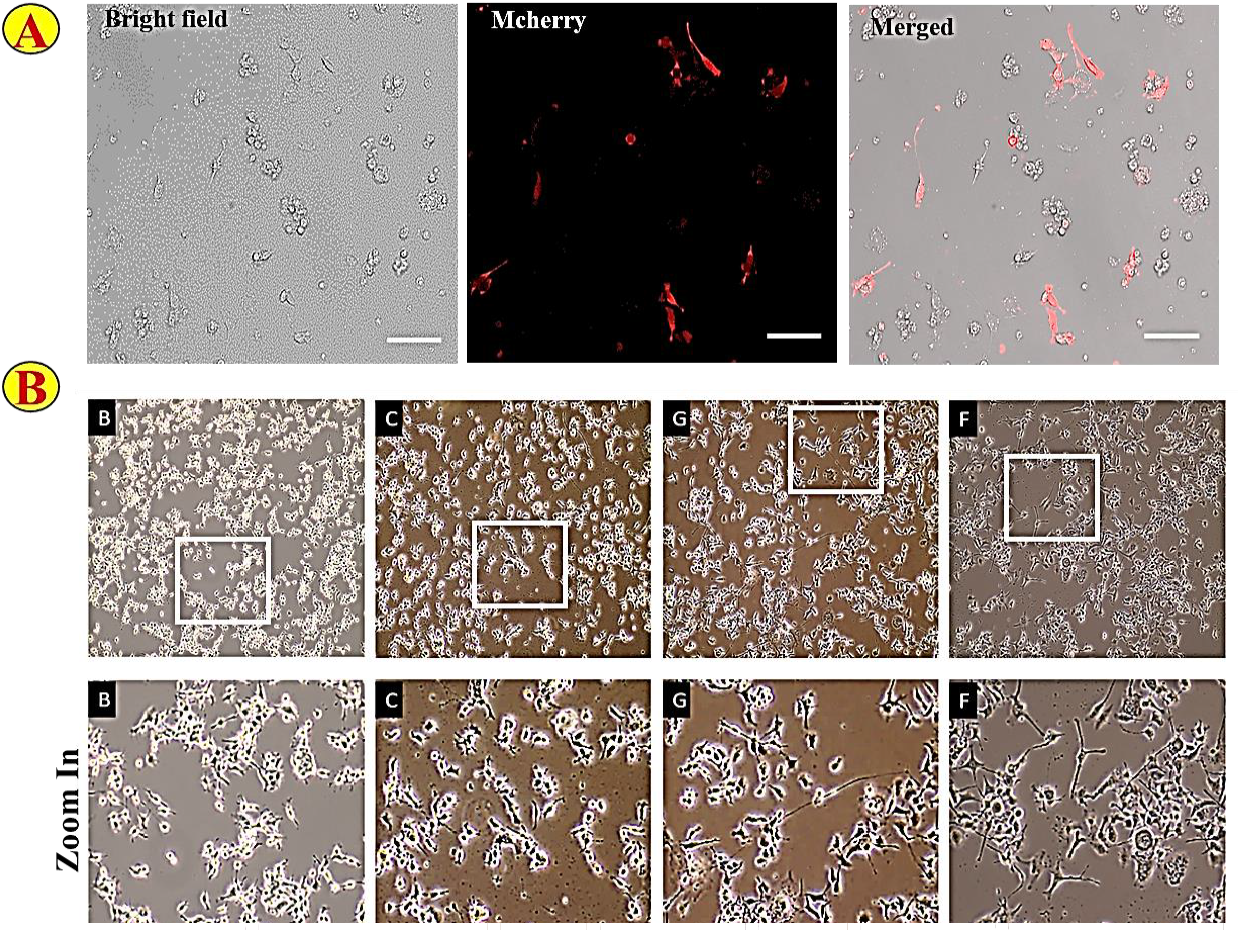
CRISPR/Cas9 Transfection Efficiency and Single Cell Clone Selection. A) N2a cells were transfected with a CRISPR-Cas9 vector containing twin guide sgRNAs, the Cas9 nuclease, and an mCherry reporter to monitor transfection. Representative images show N2a cells under brightfield (left), mCherry fluorescence (middle), and merged channels (right). B) Representative bright field images (upper panel) show the cell growth and distribution, while lower panel displays high magnification (zoom in) views revealing detailed cellular morphology and colony structure. Letters B, C, G, and F indicate the four surviving clones that were successfully maintained throughout the propagation period.

### 3.2 Molecular validation of SQSTM1/p62 knockout

To confirm successful disruption of the SQSTM1 locus, genomic DNA from wild-type (WT) cells and selected CRISPR-edited clones was analyzed by PCR using primers flanking the target region. Temperature-gradient PCR revealed altered amplification patterns in clone F compared with WT cells, consistent with genomic modification at the target site (Fig. 2A). Sequence analysis of PCR products further confirmed the presence of deletions within the SQSTM1 coding region in clone F, indicating successful CRISPR-mediated editing. Loss of p62 protein expression was verified by immunoblot analysis. WT cells showed robust p62 expression, whereas p62 protein was undetectable in clone F and markedly reduced in other edited clones (Fig. 2B). Equal protein loading was confirmed using a loading control. These results demonstrate successful knockout of SQSTM1 at both the genomic and protein levels. Based on these validations, clone F was selected for all subsequent functional analyses.

**Figure 2.**
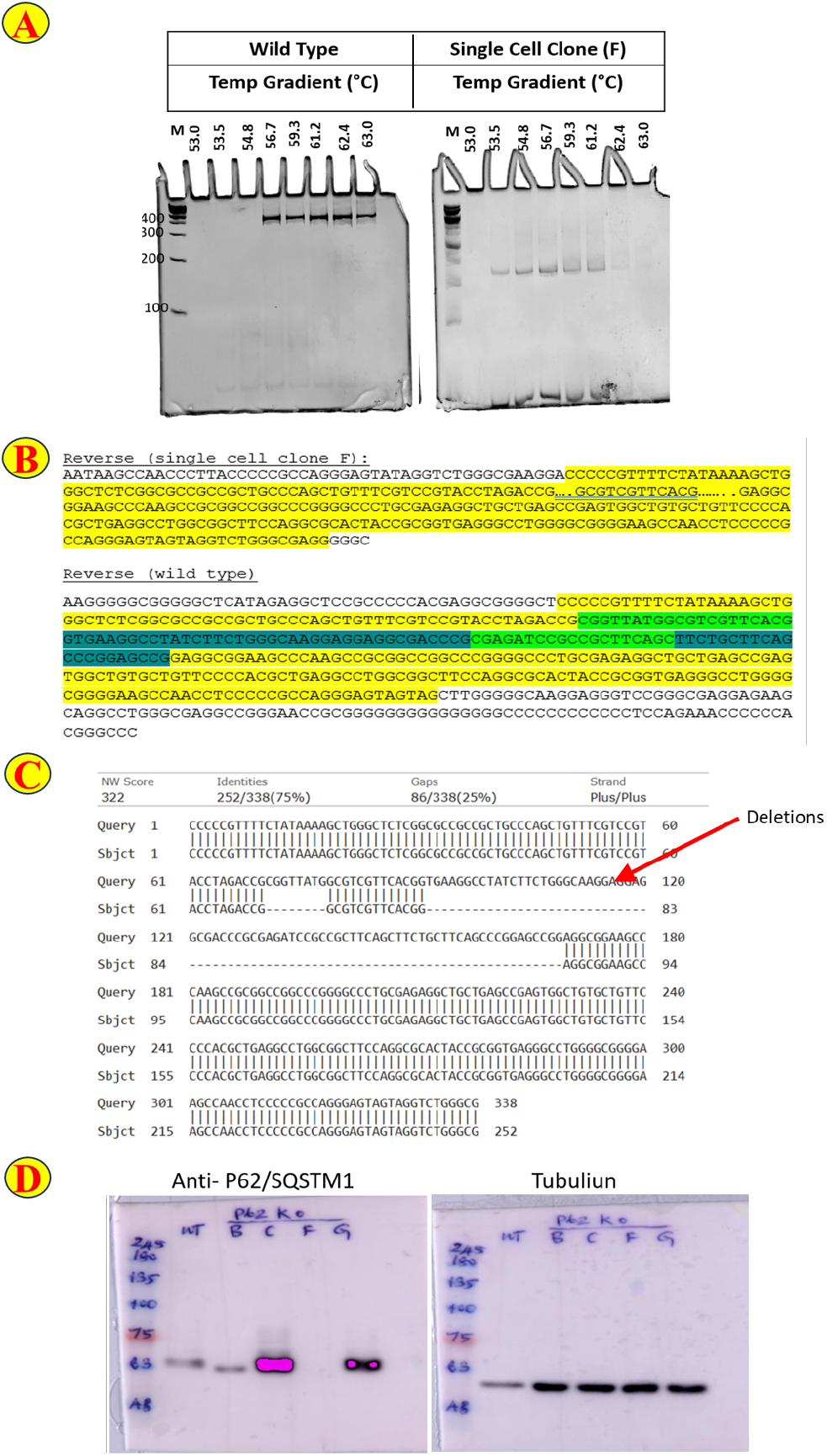
Validation of CRISPR–Cas9–mediated SQSTM1/p62 knockout clones. **(A)** Genotyping analysis of CRISPR–Cas9–edited N2a cells by temperature-gradient PCR. Genomic DNA from wild-type (WT) cells and a representative single-cell–derived clone (clone F) was amplified using primers flanking the CRISPR target site across a temperature gradient ranging from 53.0 °C to 63.0 °C. WT cells exhibited a consistent PCR product across all annealing temperatures, whereas clone F showed an altered amplification pattern, indicative of insertions or deletions (indels) at the SQSTM1 locus. Lane M denotes the DNA molecular weight marker, with fragment sizes indicated in base pairs (bp).**(B)** Sanger sequencing analysis of the PCR-amplified SQSTM1 target region from WT cells and clone F. Highlighted regions indicate sequence alterations in clone F relative to WT, confirming CRISPR–Cas9–mediated disruption of the SQSTM1 gene.**(C)** BLAST global alignment of sequencing reads from clone F against the WT SQSTM1 reference sequence. The alignment reveals substantial deletions within the targeted region, with 86 gaps out of 338 nucleotides (∼25%), further validating successful gene editing. **(D)** Immunoblot analysis of p62/SQSTM1 protein expression in WT cells and CRISPR-edited single-cell clones (B, C, F, and G). Cell lysates were resolved by SDS–PAGE and probed with anti-p62/SQSTM1 antibody (left panel). Tubulin was used as a loading control (right panel). WT cells displayed robust p62 expression, whereas clone F showed complete loss of p62 protein, confirming functional knockout. Pink-highlighted bands indicate saturating exposure during image acquisition using the Amersham Imager 600.

### 3.3 Optimization of lysosomal inhibition conditions for autophagic flux measurements

To establish optimal conditions for lysosomal inhibition and reliable measurement of autophagic flux in N2a cells, bafilomycin A1 (Baf A1) was tested across a range of concentrations and treatment durations. Cells were treated with increasing concentrations of Baf A1 (5–80 nM) for either 2 h or 4 h, and accumulation of LC3-II was assessed by immunoblotting as a readout of lysosomal blockade (Fig. 3A). Baf A1 treatment induced a dose-dependent increase in LC3-II levels at both time points, with maximal accumulation observed at concentrations ≥10 nM. Quantitative analysis revealed that LC3-II-fold change plateaued at 10–20 nM Baf A1 for both 2 h and 4 h treatments, with no substantial additional increase at higher concentrations (Fig. 3B). Based on these results, treatment with 10 nM Baf A1 for 2 h was selected for all subsequent autophagic flux experiments, as it provided robust and reproducible LC3-II accumulation while minimizing prolonged exposure

**Figure 3.**
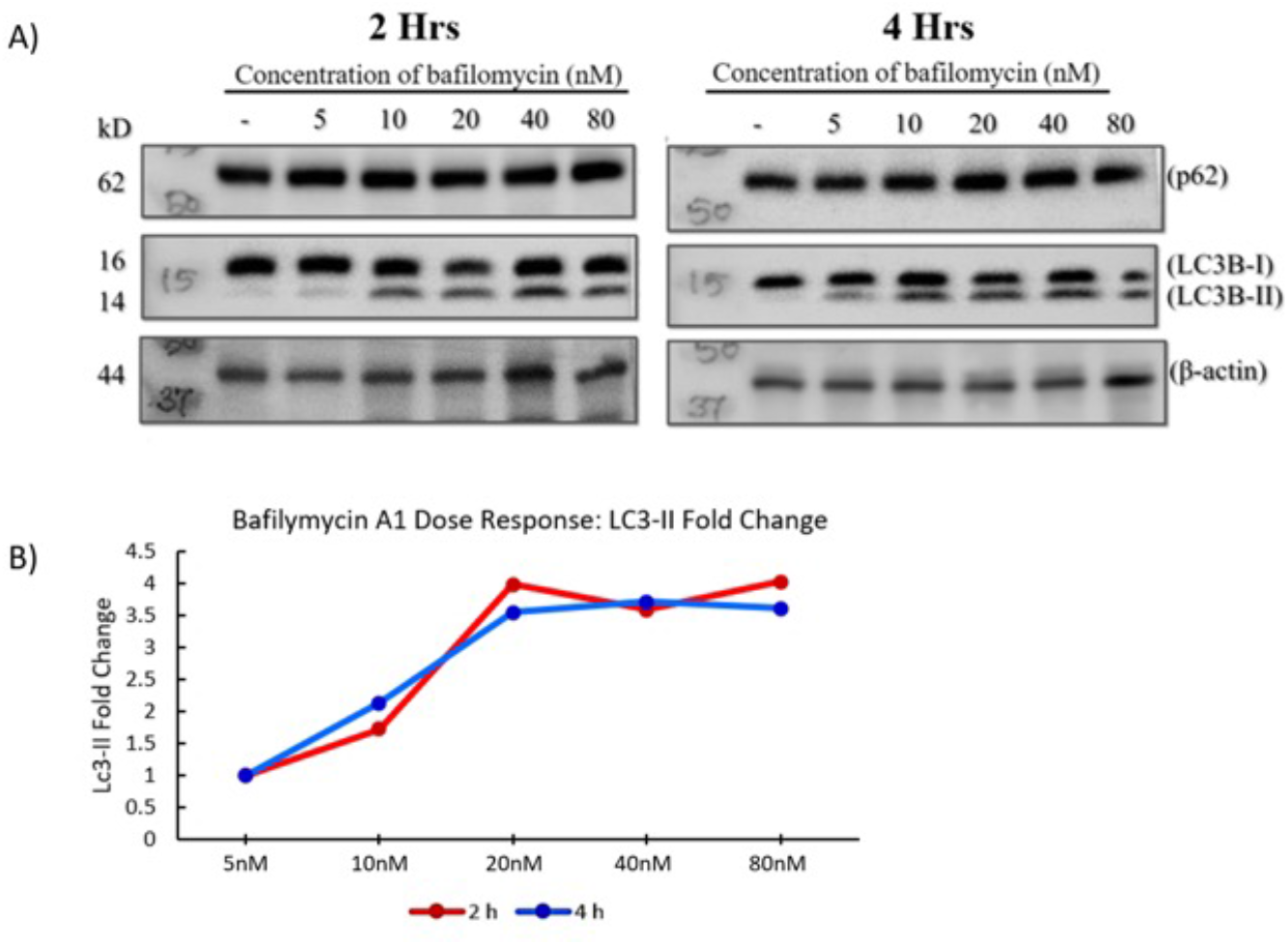
Optimization of bafilomycin A1 treatment conditions for autophagic flux analysis. **(A)** Representative immunoblots showing p62, LC3B (LC3-I and LC3-II), and β-actin levels in N2a cells treated with increasing concentrations of bafilomycin A1 (5–80 nM) for 2 h or 4 h. Accumulation of LC3-II was used as a readout of lysosomal inhibition. **(B)** Quantification of LC3-II-fold change relative to untreated controls following bafilomycin A1 treatment for 2 h (red) or 4 h (blue). LC3-II levels increased in a dose-dependent manner and plateaued at concentrations ≥10 nM for both treatment durations.

### 3.4 p62 is dispensable for basal autophagic flux in N2a cells

To determine whether p62 contributes to basal autophagy in N2a cells, WT and p62^−^/^−^ N2a cells were cultured under nutrient-replete conditions and treated with or without Baf A1. Autophagic flux was quantified by measuring the difference in LC3-II levels between Baf A1–treated and untreated cells. Under basal conditions, LC3-II accumulation in response to Baf A1 was comparable between WT and p62^−^/^−^ cells, resulting in similar autophagic flux values (Fig. 4B). These findings indicate that basal autophagic flux in N2a cells is largely preserved in the absence of p62.

**Figure 4.**
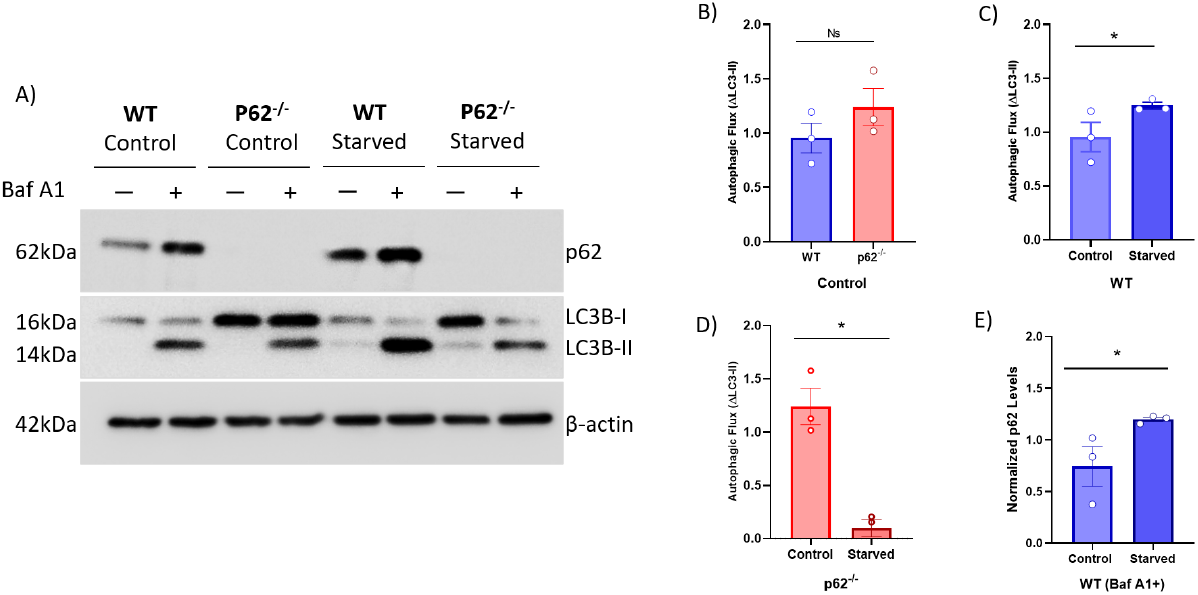
p62 is required for efficient starvation-induced autophagic flux. (A) Representative immunoblots showing p62, LC3B, and β-actin levels in wild-type (WT) and p62^−^/^−^ cells cultured under control or starvation conditions in the presence (+) or absence (−) of bafilomycin A1 (Baf A1).(B) Quantification of basal autophagic flux, calculated as the difference in LC3-II levels between Baf A1-treated and untreated WT and p62^−^/^−^ cells under control conditions.(C) Starvation-induced autophagic flux in WT cells.(D) Starvation-induced autophagic flux in p62^−^/^−^ cells.(E) Quantification of p62 protein levels in WT cells under control and starvation conditions in the presence of Baf A1, reflecting changes in p62 expression independent of lysosomal degradation. Data are presented as mean ± SEM with individual biological replicates shown. Statistical significance was assessed using unpaired two-tailed t-tests with Welch’s correction. *P < 0.05; ns, not significant.

### 3.5 p62 is required for starvation-induced autophagic flux

We next examined whether p62 is required for autophagy induction in response to nutrient deprivation. WT and p62^−^/^−^ cells were subjected to starvation using EBSS, and autophagic flux was assessed by LC3-II accumulation in the presence and absence of Baf A1. In WT cells, starvation resulted in a significant increase in LC3-II flux compared with control conditions, consistent with robust induction of autophagy upon nutrient deprivation (Fig. 4C). In contrast, p62^−^/^−^ cells failed to exhibit a comparable increase in LC3-II flux upon starvation and instead showed a marked reduction relative to their basal condition (Fig. 4D). These results demonstrate that p62 is required for an efficient starvation-induced autophagic response in N2a cells.

### 3.6 Starvation induces p62 expression independent of lysosomal degradation

Given the observed defect in starvation-induced autophagy in p62-deficient cells, we next examined how p62 protein levels respond to nutrient deprivation in WT cells. To assess changes in p62 expression independent of lysosomal degradation, p62 levels were quantified in WT cells treated with Baf A1 under control and starvation conditions. Starvation resulted in increased p62 protein levels in the presence of Baf A1 (Fig. 4E), indicating that nutrient deprivation induces p62 expression rather than causing accumulation due to impaired degradation. This finding supports a model in which p62 is transcriptionally or translationally upregulated during starvation while remaining subject to autophagic turnover.

## 4. Discussion

Autophagy is a fundamental cellular process that supports homeostasis and adaptation to metabolic stress. The selective autophagy receptor p62/SQSTM1 has been implicated in multiple aspects of autophagy regulation, including cargo recognition, signaling integration, and stress adaptation. However, its precise contribution to basal versus stress-induced autophagic flux remains context dependent and incompletely defined. In this study, using a neuroblastoma (N2a) cell–based neuronal precursor–like model, we demonstrate that p62 is dispensable for basal autophagic flux but is required for an efficient autophagic response to nutrient deprivation.

Under nutrient-replete conditions, loss of p62 did not significantly alter LC3-II flux in N2a cells, indicating that basal autophagy can proceed independently of p62 in this neuronal cell model. This finding is consistent with the notion that constitutive autophagy is primarily maintained by the core autophagy machinery, while selective autophagy receptors may play a more prominent role under conditions of increased cellular stress or cargo demand. Previous studies have similarly reported that p62 deficiency does not universally impair basal autophagy, supporting the idea that p62 function is highly context dependent rather than obligatory for all autophagic activity (you may cite general p62/autophagy reviews here).

In contrast, starvation robustly induced autophagic flux in wild-type N2a cells but failed to do so in p62-deficient cells, revealing a specific requirement for p62 during nutrient stress. Because autophagic flux was assessed using lysosomal inhibition, the observed defect reflects impaired autophagy induction or maintenance rather than altered lysosomal degradation. These results suggest that p62 contributes to the cellular capacity to upregulate autophagy in response to metabolic stress, potentially by facilitating selective cargo handling, scaffolding autophagosome assembly, or sustaining autophagic signaling under starvation conditions. Similar stress-dependent roles for p62 have been reported in other systems, where p62 supports adaptive autophagy rather than basal turnover.

Consistent with this stress-responsive role, p62 protein levels increased in wild-type cells during starvation when lysosomal degradation was blocked. Importantly, p62 continued to accumulate upon bafilomycin A1 treatment, indicating that elevated p62 levels reflect enhanced expression rather than defective autophagic clearance. p62 is known to be transcriptionally regulated by stress-responsive pathways, including NRF2- and ATF4-dependent programs, which are activated during nutrient deprivation. Increased p62 expression under starvation may therefore serve to expand selective autophagy capacity or reinforce autophagosome dynamics during periods of elevated autophagic demand.

Taken together, our findings support a model in which p62 plays a context-dependent role in autophagy regulation in a neuronal cell model. While basal autophagy proceeds largely independently of p62, p62 becomes essential for mounting an effective autophagic response to nutrient stress. Given that N2a cells are widely used as a neuronal precursor–like model, these observations may have implications for understanding how autophagy is adapted during metabolic stress in neuronal contexts. Future studies using differentiated neurons or in vivo models will be required to determine how p62-dependent autophagy regulation contributes to neuronal stress resilience under physiological and pathological conditions.

## Notes

### Competing Interest Statement

The authors have declared no competing interest.

### Funding

- Department of Science and Technology- SERB-CRG
- DAAD
- UGC - JRF

## References

Agudo-Canalejo, J., Schultz, S. W., Chino, H., Migliano, S. M., Saito, C., Koyama-Honda, I., Stenmark, H., Brech, A., May, A. I., Mizushima, N., & Knorr, R. L. (2021). Wetting regulates autophagy of phase-separated compartments and the cytosol. Nature, 591(7848), 142–146. 10.1038/S41586-020-2992-3,

Bjørkøy, G., Lamark, T., Brech, A., Outzen, H., Perander, M., Øvervatn, A., Stenmark, H., & Johansen, T. (2005). p62/SQSTM1 forms protein aggregates degraded by autophagy and has a protective effect on huntingtin-induced cell death. Journal of Cell Biology, 171(4), 603–614. 10.1083/JCB.200507002,

Fan, W., Tang, Z., Chen, D., Moughon, D., Ding, X., Chen, S., Zhu, M., & Zhong, Q. (2010). Keap1 facilitates p62-mediated ubiquitin aggregate clearance via autophagy. Autophagy, 6(5), 614–621. 10.4161/AUTO.6.5.12189,

Geisler, S., Holmström, K. M., Skujat, D., Fiesel, F. C., Rothfuss, O. C., Kahle, P. J., & Springer, W. (2010). PINK1/Parkin-mediated mitophagy is dependent on VDAC1 and p62/SQSTM1. Nature Cell Biology, 12(2), 119–131. 10.1038/NCB2012,

Ichimura, Y., Kumanomidou, T., Sou, Y. S., Mizushima, T., Ezaki, J., Ueno, T., Kominami, E., Yamane, T., Tanaka, K., & Komatsu, M. (2008). Structural basis for sorting mechanism of p62 in selective autophagy. Journal of Biological Chemistry, 283(33), 22847–22857. 10.1074/jbc.M802182200

Kageyama, S., Gudmundsson, S. R., Sou, Y. S., Ichimura, Y., Tamura, N., Kazuno, S., Ueno, T., Miura, Y., Noshiro, D., Abe, M., Mizushima, T., Miura, N., Okuda, S., Motohashi, H., Lee, J. A., Sakimura, K., Ohe, T., Noda, N. N., Waguri, S., … Komatsu, M. (2021). p62/SQSTM1-droplet serves as a platform for autophagosome formation and anti-oxidative stress response. Nature Communications, 12(1). 10.1038/S41467-020-20185-1,

Katsuragi, Y., Ichimura, Y., & Komatsu, M. (2015). P62/SQSTM1 functions as a signaling hub and an autophagy adaptor. In FEBS Journal (Vol. 282, Issue 24, pp. 4672–4678). Blackwell Publishing Ltd. 10.1111/febs.13540

Klionsky, D. J., Abdelmohsen, K., Abe, A., Abedin, M. J., Abeliovich, H., Arozena, A. A., Adachi, H., Adams, C. M., Adams, P. D., Adeli, K., Adhihetty, P. J., Adler, S. G., Agam, G., Agarwal, R., Aghi, M. K., Agnello, M., Agostinis, P., Aguilar, P. V., Aguirre-Ghiso, J., … Zughaier, S. M. (2016). Guidelines for the use and interpretation of assays for monitoring autophagy (3rd edition). Autophagy, 12(1), 1. 10.1080/15548627.2015.1100356

Komatsu, M. (2022). p62 bodies: Phase separation, NRF2 activation, and selective autophagic degradation. In IUBMB Life (Vol. 74, Issue 12, pp. 1200–1208). John Wiley and Sons Inc. 10.1002/iub.2689

Komatsu, M., Waguri, S., Koike, M., Sou, Y. shin, Ueno, T., Hara, T., Mizushima, N., Iwata, J. ichi, Ezaki, J., Murata, S., Hamazaki, J., Nishito, Y., Iemura, S. ichiro, Natsume, T., Yanagawa, T., Uwayama, J., Warabi, E., Yoshida, H., Ishii, T., … Tanaka, K. (2007). Homeostatic Levels of p62 Control Cytoplasmic Inclusion Body Formation in Autophagy-Deficient Mice. Cell, 131(6), 1149–1163. 10.1016/j.cell.2007.10.035

Mizushima, N., Yoshimori, T., & Ohsumi, Y. (2011). The role of atg proteins in autophagosome formation. Annual Review of Cell and Developmental Biology, 27, 107–132. 10.1146/ANNUREV-CELLBIO-092910-154005,

Nakamura, M., Verboon, J. M., Prentiss, C. L., & Parkhurst, S. M. (2020). The kinesin-like protein Pavarotti functions noncanonically to regulate actin dynamics. Journal of Cell Biology, 219(9). 10.1083/JCB.201912144,

Nixon, R. A. (2013). The role of autophagy in neurodegenerative disease. Nature Medicine, 19(8), 983–997. 10.1038/NM.3232;SUBJMETA=1283,1689,378,631;KWRD=ALZHEIMER

Pankiv, S., Clausen, T. H., Lamark, T., Brech, A., Bruun, J. A., Outzen, H., Øvervatn, A., Bjørkøy, G., & Johansen, T. (2007). p62/SQSTM1 binds directly to Atg8/LC3 to facilitate degradation of ubiquitinated protein aggregates by autophagy*[S]. Journal of Biological Chemistry, 282(33), 24131–24145. 10.1074/jbc.M702824200

Ravenhill, B. J., Boyle, K. B., von Muhlinen, N., Ellison, C. J., Masson, G. R., Otten, E. G., Foeglein, A., Williams, R., & Randow, F. (2019). The Cargo Receptor NDP52 Initiates Selective Autophagy by Recruiting the ULK Complex to Cytosol-Invading Bacteria. Molecular Cell, 74(2), 320-329.e6. 10.1016/j.molcel.2019.01.041

Sánchez-Martin, P., & Komatsu, M. (2018). p62/SQSTM1 – Steering the cell through health and disease. Journal of Cell Science, 131(21). 10.1242/JCS.222836/77085

Turco, E., Witt, M., Abert, C., Bock-Bierbaum, T., Su, M. Y., Trapannone, R., Sztacho, M., Danieli, A., Shi, X., Zaffagnini, G., Gamper, A., Schuschnig, M., Fracchiolla, D., Bernklau, D., Romanov, J., Hartl, M., Hurley, J. H., Daumke, O., & Martens, S. (2019). FIP200 Claw Domain Binding to p62 Promotes Autophagosome Formation at Ubiquitin Condensates. Molecular Cell, 74(2), 330. 10.1016/J.MOLCEL.2019.01.035

Vargas, J. N. S., Wang, C., Bunker, E., Hao, L., Maric, D., Schiavo, G., Randow, F., & Youle, R. J. (2019). Spatiotemporal Control of ULK1 Activation by NDP52 and TBK1 during Selective Autophagy. Molecular Cell, 74(2), 347-362.e6. 10.1016/j.molcel.2019.02.010

